# Hedgehog Activity Gradient in Combination with Transcription Network Confers Multiple Hypothalamic Identities

**DOI:** 10.1101/2022.08.23.505035

**Authors:** Maho Yamamoto, Agnes Ong Lee Chen, Takuma Shinozuka, Manabu Shirai, Noriaki Sasai

## Abstract

During development, the hypothalamus emerges from the ventral diencephalon of the neural tube and is regionalised into several distinct functional domains. Each domain is characterised by different combinations of transcription factors, the expression of which is regulated by signalling molecules and downstream transcriptional networks. Transcription factors, including Nkx2.1, Nkx2.2, Pax6 and Rx, are expressed in the presumptive hypothalamus and its surrounding regions from an early developmental stage and play critical roles in the development of these areas. However, the regulation of transcription factor expression and the details of the transcriptional network among them have not been fully elucidated.

As early hypothalamus development takes place in the ventral region of the forebrain where Sonic Hedgehog (Shh) is expressed, we focused on the relationship between Shh and its downstream transcription factors and investigated the transcriptional regulation along the dorsal-ventral axis. Using a reporter mouse line, *in vitro* neural differentiation of mouse embryonic stem cells and gene overexpression in chick embryos, we found that Pax6, Nkx2.1 and Nkx2.2 are regulated epistatically by different Shh signal intensities. Nkx2.1 and Nkx2.2 mutually repress each other; however, they induce each other in a non-cell-autonomous manner. Moreover, Rx resides upstream of all these transcription factors and determines the location of the hypothalamic region along the dorsal-ventral and anterior-posterior regulations. Finally, we found that the Shh signal demarcates the diencephalic region from the retinal area. Our findings suggest that Shh signalling, and its downstream transcription network, are required for hypothalamic regionalisation and establishment of diencephalic cell fate.

## Introduction

The hypothalamus acts as an endocrine hub and plays an important role in the homeostasis of autonomous activities and behavioural functions (Biran et al., 2015). During development, the hypothalamus emerges as a domain in the ventral portion of the diencephalon, with the epithalamus and thalamus as dorsal segments (Diaz and Puelles, 2020; Puelles and Rubenstein, 2003).

During development, the hypothalamus is further divided into several subdomains along the dorsal-ventral (D-V) and rostral-caudal (R-C) axes (Diaz and Puelles, 2020; Kim et al., 2020; Shimogori et al., 2010) corresponding to distinct functions. Each domain comprises different sets of transcription factors, many of which are homeobox-type (Bedont et al., 2015; Kim et al., 2020; Shimogori et al., 2010) and acquire unique characteristics (Diaz and Puelles, 2020). Representative transcription factors expressed at the earliest stages of hypothalamic development are Nkx2.1 (Orquera et al., 2019; Shimamura and Rubenstein, 1997; Sussel et al., 1999), Nkx2.2 (Briscoe et al., 1999; Orquera et al., 2016) and Rx (Furukawa et al., 1997; Mathers et al., 1997). The expression profiles of these transcription factors differ slightly (Orquera et al., 2016) and are essential for establishing a detailed hypothalamic pattern (Wataya et al., 2008).

The emergence of these subdomains is dependent on secreted proteins, such as Sonic Hedgehog (Shh) (Diaz and Puelles, 2020; Kim et al., 2020; Lupo et al., 2006; Manning et al., 2006; Ohyama et al., 2008; Shimamura and Rubenstein, 1997), Bone Morphogenetic Protein (BMP) (Dale et al., 1997; Manning et al., 2006; Ohyama et al., 2008), Fibroblast Growth Factor (FGF) (Tsai et al., 2011) and Wnt (Harrison-Uy and Pleasure, 2012; Newman et al., 2018). These signalling molecules generate gradients and provide positional information on progenitor cells along the anterior-posterior (A-P) and D-V axes in the brain. Transcription factors respond to these signals depending on their concentration and combinations, and the subdomains are positioned in a spatiotemporally precise manner.

Among them, Shh is expressed in the ventral neural tube and its underlying prechordal plate from the early stages of the forebrain (Diaz and Puelles, 2020; Lupo et al., 2006; Ohyama et al., 2008; Szabo et al., 2009; Traiffort et al., 2016) and has been shown to be critical for the development and patterning of ventral forebrain regions (Bertrand and Dahmane, 2006; Chiang et al., 1996; Ericson et al., 1995);Blaess, 2014 #38}. At the cellular level, Shh activates the transcription factor Gli, which translocates from the cell membrane to the nucleus and induces the expression of target genes. The flanking regions *of Nkx2.1* (Shimamura and Rubenstein, 1997) and *Nkx2.2* (Briscoe et al., 1999) contain Gli-binding sites and are directly activated by Shh (Kutejova et al., 2016; Nishi et al., 2015; Vokes et al., 2007). Conversely, the expression of the homeodomain protein *Pax6* is downregulated (Briscoe et al., 2000); this repression is mediated by the direct binding of Nkx2.2 in the gene regulatory region (Kutejova et al., 2016). The combination of positional information provided by Shh and the downstream transcriptional network formed by the target genes establishes a variety of subdomains within the neural tube with distinct boundaries, precise order, and an accurate number of cells.

Nkx2.1, Nkx2.2 and Rx are expressed in the rostral hypothalamus (Bedont et al., 2015; Wataya et al., 2008), whereas Pax6 is expressed in the dorsal hypothalamus (Corman et al., 2018; Kioussi et al., 1999) during the early stages of forebrain development. Rx is broadly expressed in the forebrain (Orquera et al., 2016), and Nkx2.1-positive/Nkx2.2-negative cells further form the arcuate nucleus (ARC) neurons, whereas double-positive cells for Nkx2.1 and Nkx2.2 differentiate into the ventromedial hypothalamus (VMH) (Bedont et al., 2015). However, the transcriptional network between these factors during the initial developmental stages is not fully understood.

In the present study, we first used reporter mice to monitor Hedgehog (Hh) signalling (Balaskas et al., 2012) and describe the relationship between dynamic Gli activity and hypothalamic transcription factor profiles. Next, we used an *in vitro* neural differentiation system to differentiate mouse embryonic stem (ES) cells into hypothalamic neural cells to uncover the mutual dependencies of transcription factors during this process. We also exploited gene overexpression in chick embryos to address the effects of transcription factor overexpression. Our data highlight the significance of the combination of permissive and instructive functions in hypothalamic gene expression.

## Results

### Gli activity coincides with transcription factor expression in developing forebrain regions

To address the relationship between Hh signalling and transcription factor expression characteristics of diencephalic patterning, we used *Tg (GBS-Gfp)* mice, where the spatial and temporal distributions of Gli activity were monitored in a timely manner (Balaskas et al., 2012).

We obtained timed pregnant mice of *Tg (GBS-Gfp)*, and extracted embryos at embryonic days of 8.5 (e8.5) to e10.5, when forebrain patterning was progressive. Gli activity, reported through GFP expression, was detected in the ventral forebrain at e8.5 (Fig. 1A) and at e9.5 in the intermediate (yellow arrowhead; Fig. 1B) and ventral regions of the diencephalon (white bracket; Fig. 1B). At e10.5, weak Gli activity was observed in the presumptive medial ganglionic eminence (MGE; yellow arrowhead; Fig. 1C) and strong Gli activity was observed in the dorsal hypothalamus (white bracket: Fig. 1C). The retinal area, which evaginates from the neural tube and forms a part of the eye, did not exhibit Gli activity (open arrowheads; Fig. 1C). These data suggested that Gli was active in the ventral region of the forebrain during development (Fig. 1A–C).

**Fig. 1.**
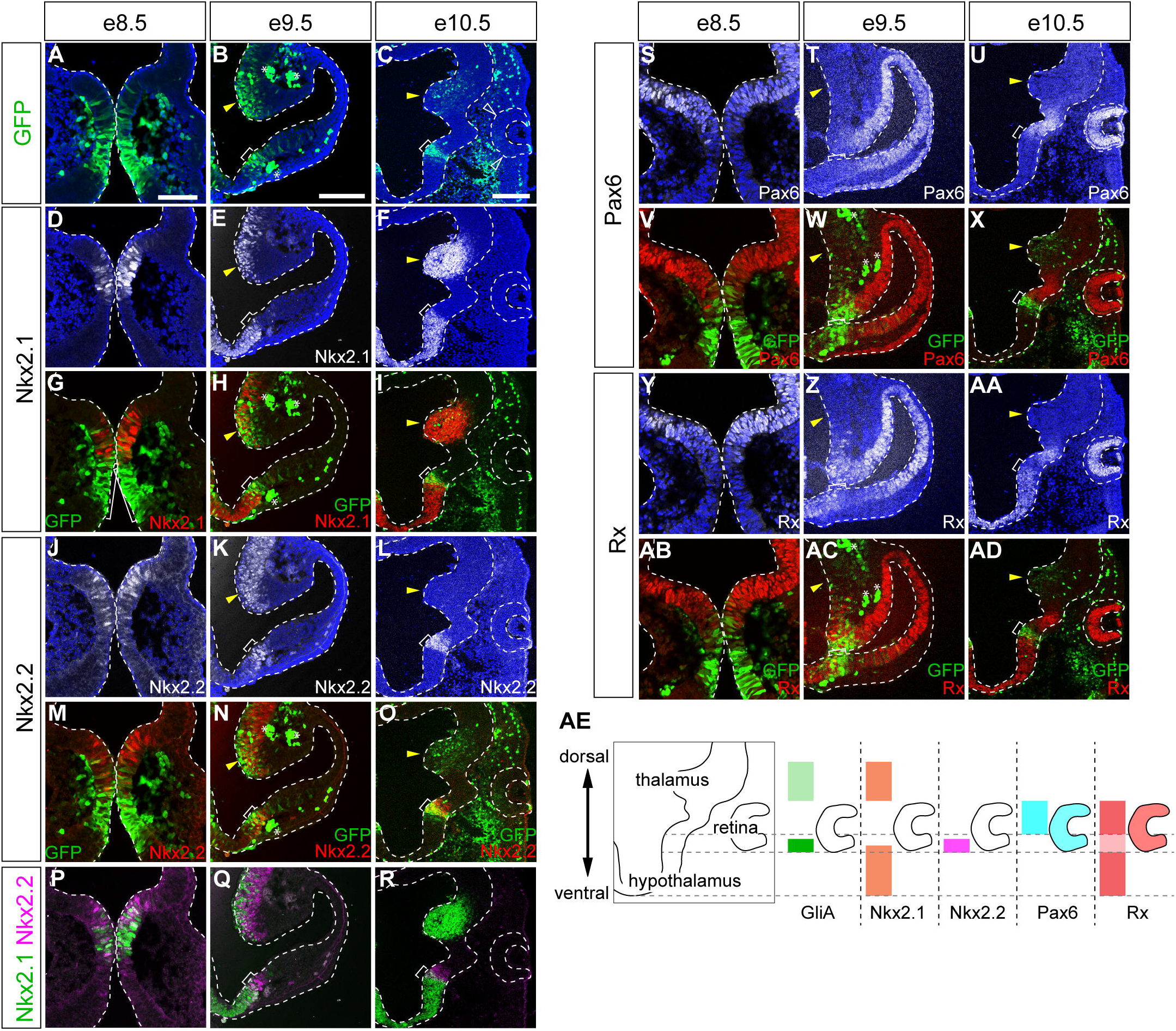
Expression of ventral diencephalic transcription factors correlate with Gli activity. (A–C) Distribution of Gli activity (GFP expression, green) was monitored at e8.5 (A), e9.5 (B) and e10.5 (C), co-staining with DAPI (blue; A–C). (D–AD) Expression of Nkx2.1 (white in D–F; red in G-I), Nkx2.2 (white in J–L; red in M–O), Pax6 (white in S–U; red in V–X) and Rx (white in Y–AA; red in AB–AD) was analysed using GFP immunofluorescence (reflecting Gli activity; G–I,M–O,V–X,AB– AD). Nkx2.1 and Nkx2.2 expression are merged (P–R; green for Nkx2.1 and purple for Nkx2.2). Non-specific GFP staining is indicated with asterisks. Areas with high GFP expression are indicated by brackets, presumptive MGE with yellow arrowheads. Four embryos were analysed for each stage. Scale bars, 50 μm (A for A,D,G,J,M,P,S,V,Y,AB), 100 μm (B for B,E,H,K,N,Q,T,W,Z,AC), and 200 μm (C for C,F,I,L,O,R,U,X,AA,AD). (AE) Schematic of the distributions of either Gli activity or expression of Nkx2.1, Nkx2.2, Pax6 and Rx along the dorsal-ventral axis, focusing on hypothalamus, thalamus and retina.

Next, we investigated the correlation between the expression of transcription factors involved in forebrain development and patterning with temporal Gli activity. We first focused on the transcription factors Nkx2.1 and Nkx2.2 as they are directly induced by Shh (Briscoe et al., 1999; Dessaud et al., 2007; Shimamura and Rubenstein, 1997).

At e8.5, Nkx2.1 and Nkx2.2 expression was found in the area where Gli activity was evident (Fig. 1D,G,J,M). At e9.5, Nkx2.1 and Nkx2.2 expression was detected in the intermediate region (yellow arrowheads; Fig. 1E,H,K,N) and in the ventral diencephalic area. Nkx2.1 expression reached the ventral midline (Fig. 1E,H), whereas Nkx2.2 expression was limited to the area with high Gli activity (Fig. 1K,N). At e10.5, Nkx2.1 was expressed widely in the hypothalamic region (Fig. 1F,I). Nkx2.2 expression was restricted to the area with high Gli activity (Fig. 1L,O), and the expression in the presumptive MGE almost disappeared (yellow arrowheads; Fig. 1L,O).

These findings support the idea that Nkx2.1 and Nkx2.2 expression is largely dependent on Hh signalling; Nkx2.1- and Nkx2.2-expressing cells emerge in the area where GFP expression is evident (Fig. 1D–O). However, their expression patterns were different. Nkx2.1 and Nkx2.2 expression were complementary to each other (Fig. 1P–R) in the area where Gli activity was low; however, several double positive cells for Nkx2.1 and Nkx2.2 were found in the area with high Gli activity (Fig. 1P–R), suggesting that there is Gli-dependent regulation between the two transcription factors. However, in the areas where Gli activity was high, Nkx2.1 and Nkx2.2 were co-expressed (Fig. 1Q,R; brackets), which further suggested that the repressive effect is dependent on Hh activity.

We also investigated the expression of other transcription factors that are involved in forebrain patterning. At e8.5, Pax6 expression was extensive in the forebrain region but was complementary to the area where Gli activity was detected (Fig. 1S,V). This trend persisted throughout all the embryonic days analysed, suggesting that Pax6 is expressed where Gli activity is repressed. Moreover, Pax6 expression was observed in the presumptive retinal area at e9.5 and e10.5 (Fig. 1T,W,U,X).

For another transcription factor, Rx, the expression pattern in the forebrain was nearly the same as that of Pax6 at e8.5 (Fig. 1Y,AB) and e9.5 (Fig. 1Z,AC). At e10.5 (Fig. 1AA,AD), the expression was detected both in the ventral diencephalic (Nkx2.1-positive; induced by Shh) and presumptive retinal areas (Pax6-positive; repressed by Shh), suggesting that Rx expression is less affected by the Hh signal.

These data showed that the expression of hypothalamic transcription factors correlated with Gli activity (Fig. 1AE). However, their dependency on Shh signalling is different.

### Nkx2.1 and Nkx2.2 are induced differentially by Hh signal

We next sought to recapitulate the hypothalamic identities *in vitro* using a neural differentiation system with murine ES cells. Ventral forebrain identities have been obtained through differentiation of ES cells with growth factor-free chemically defined medium (gfCDM) (Wataya et al., 2008). This neural differentiation system addresses the direct effects of inputs on gene expression. In this analysis, we used a three-dimensional differentiation culture to form colonies in a 96-well non-absorbable plate for seven days (see Materials and Methods for details) with treatment of Smoothened agonist (SAG), a chemical mimicking Hh activity, on day 3.

The sections of the colonies were analysed using immunofluorescence. In the absence of SAG, colonies did not show any expression of Nkx2.1 or Nkx2.2 (Fig. 2A,B,B’,Q,R), and were dominated by cells positive for Pax6 (Fig. 2C,T), suggesting that ES cells differentiated into neural cells but did not acquire hypothalamic identity. In contrast, Nkx2.1- and Nkx2.2-positive cells were found upon differentiation with 5 nM SAG, with a larger number of Nkx2.1-positive cells (Fig. 2E,F,F’,Q,R), indicating that SAG provided hypothalamic identity to the differentiating neural cells. Moreover, most cells expressed either Nkx2.1 or Nkx2.2, and only a few cells were double-positive for Nkx2.1 and Nkx2.2 (Fig. 2E,F,F’,Q,R,S). In contrast, higher concentrations of SAG (i.e.; 50 nM and 500 nM) provided more cells positive for Nkx2.1 and Nkx.2.2 (Fig. 2I,J,J’,Q,R,S), with most cells being positive for Nkx2.1 (Fig. 1I,M). Notably, nearly all cells positive for Nkx2.2 were also positive for Nkx2.1 with 500 nM SAG (Fig. 2M,N,N’,Q,R,S). These observations suggest that different types of hypothalamic neural progenitor cells can be generated using this ES cell differentiation system by introducing distinct Hh activity.

**Fig. 2.**
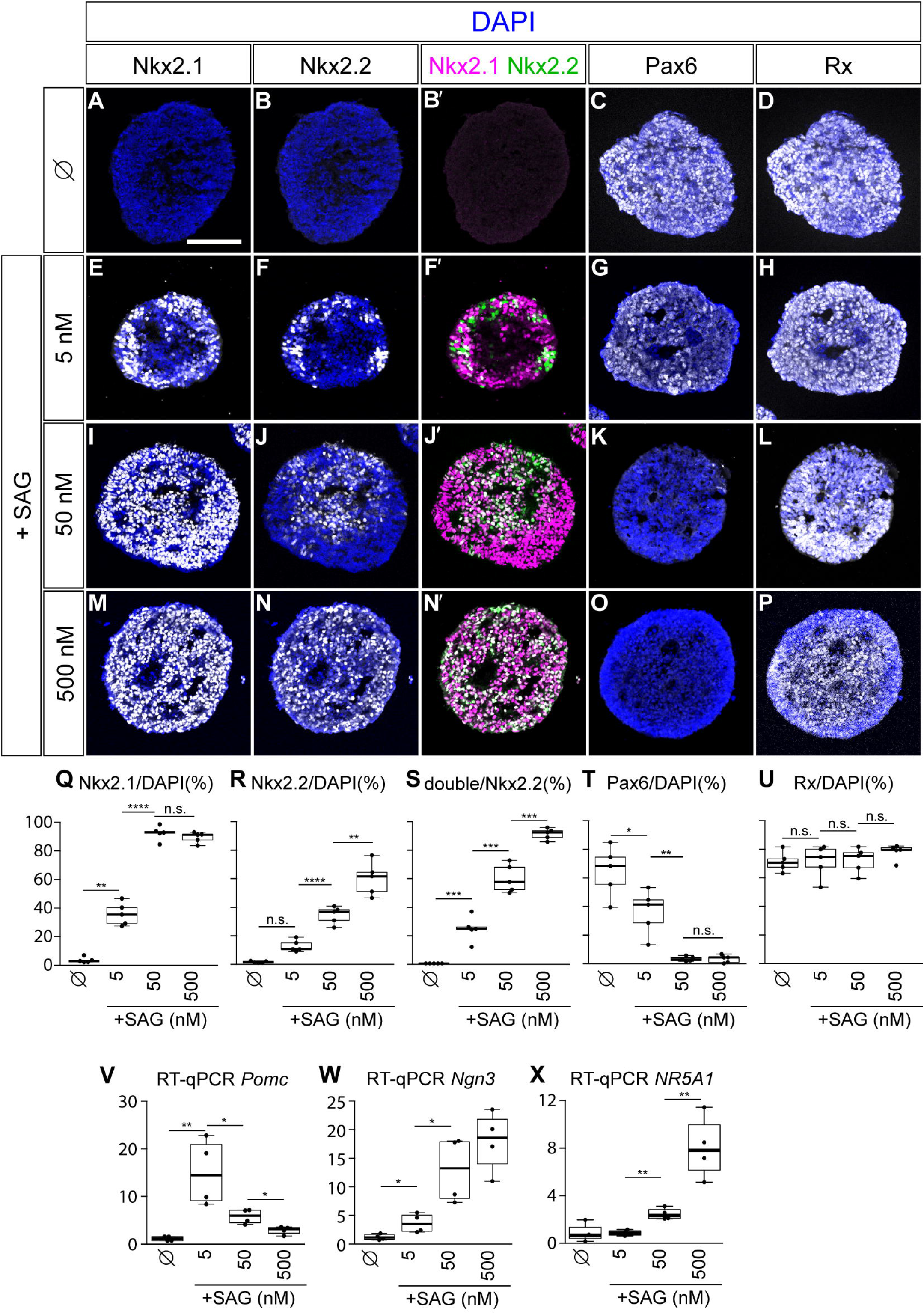
Differential Hh levels induce distinct hypothalamic domains. (A–P) ES cells were differentiated with gfCDM in the absence (A–D) or presence of 5 nM (E–H), 50 nM (I–L) or 500 nM (M–P) SAG, and the expression Nkx2.1 (A,B’,E,F’,I,J’,M,N’), Nkx2.2 (B,B’,F,F’,J,J’,N,N’), Pax6 (C,G,K,O) and Rx (D,H,L,P) was analysed using immunofluorescence. Scale bar, 50 μm (A for A–P). (Q–U) Quantification of cells positive for Nkx2.1 (Q), Nkx2.2 (R), Pax6 (T) and Rx (U) over DAPI, and double positive for Nkx2.1/Nkx2.2 over Nkx2.2 (S). (V-X) Relative expression levels of *Pomc* (V), *Ngn3* (W) and *NR5A1* (X) in the cells differentiated for 12 days with different concentrations of SAG.

In contrast to the increase in the number of Nkx2.1- and Nkx2.2-positive cells, the number of Pax6-positive cells decreased with increasing concentrations of SAG (Fig. 2C,G,K,O,T), suggesting that Hh signalling repressed Pax6 expression. In contrast to the downregulation of Pax6, the number of Rx-expressing cells appeared to be constant with increasing concentrations of SAG, suggesting that the regulation of Rx expression is independent of SAG (Fig. 2D,H,L,P,U).

Therefore, we quantified *Rx* expression levels by reverse transcription and quantitative polymerase chain reaction (RT-qPCR) in the cells differentiated using 0, 5, 50, and 500 nM SAG. We found that the expression levels tended to decrease with increasing concentrations of SAG (Supplementary Fig. S1). Together, these findings suggest that while the number of Rx-positive cells was largely unaffected by SAG, the expression level appeared to be regulated by SAG, and Rx functions in a quantitative manner.

We further attempted to address the temporal changes in the number of cells positive for the hypothalamic transcription factors, and quantified the Nkx2.1- and Nkx2.2-positive cells upon treatment with 500 nM of SAG (treated day 3 onward) from days 4–9. As a result, Nkx2.1-positive cells began to appear as early as day 4 in a SAG-dependent manner and were evident on day 5 (Supplementary Fig. S2A; black arrowhead). In contrast, Nkx2.2 expression was found later (Supplementary Fig. S2B, black arrowhead). The number of Nkx2.1- and/or Nkx2.2-positive cells was clearly higher upon SAG treatment at all time points (Supplementary Fig. S2A,B). Regarding the Pax6-expressing cells, the number of positive cells transiently increased on day 5, suggesting that Pax6 expression is induced at the initial step of the neural differentiation at either conditions with presence or absence of SAG. However, the expression decreased quickly with 500 nM SAG, while an increasing number of cells was found in the absence of SAG (Supplementary Fig. S2C). In contrast to the above three genes of Nkx2.1, Nkx2.2 and Pax6, whose expression was dependent on SAG treatment, Rx expression was already found at day 4 regardless of the presence of SAG (green arrowhead; Supplementary Fig. S2D) and continued to increase until day 7 (Supplementary Fig. S2D). These observations suggest that *Rx* resides epistatically upstream of *Nkx2.1*, *Nkx2.2* and *Pax6*, and that its expression is upregulated during neural differentiation less dependently on Hh signalling. Thus, each hypothalamic transcription factor is also differentially regulated temporally.

Finally, we investigated whether each type of hypothalamic progenitor cell further differentiated into its corresponding neurons. To investigate this possibility, we performed prolonged differentiation with different concentrations of SAG until day 12, and analysed the expression of the genes that are expressed in specific subdomains within the hypothalamus.

As a result, a significant upregulation of *Proopiomelanocortin* (*Pomc*), which is expressed in the ARC (Bedont et al., 2015), was found with 5 nM and 50 nM SAG (Fig. 2V) as analysed by RT-qPCR, while fewer positive cells were found without SAG and with 500 nM SAG (Fig. 2V). In contrast, *Ngn3* (Aslanpour et al., 2020) and *NR5A1* (also known as *SF-1*) (Kurrasch et al., 2007), which are characteristics of the ventromedial hypothalamic (VMH) neurons derived from double-positive cells for Nkx2.1 and Nkx2.2, were markedly upregulated with 500 nM SAG (Fig. 2W,X). These results are consistent with the progenitor gene expression at day 7; Nkx2.1 at the low SAG concentration (Fig. 2E) and both Nkx2.1 and Nkx2.2 at the high SAG concentration (Fig. 2M,N).

Altogether, these findings suggest that the graded Hh signal induces distinct hypothalamic neurons differentiated from different types of progenitor cells.

### *Nkx2.1* resides epistatically upstream of *Nkx2.2* and is required for repression of Pax6

Next, we investigated the mutual regulation between Nkx2.1 and Nkx2.2. Upon treatment with low concentrations of SAG, most cells were either Nkx2.1- or Nkx2.2-positive, and double-positive cells did not appear frequently (Fig. 2E,F,F’,S). In contrast, at higher concentrations of SAG, almost all Nkx2.2-positive cells were positive for Nkx2.1 (Fig. 2M,N,N’,S). This suggests that Nkx2.1 and Nkx2.2 are mutually repressive and that higher Hh/Gli activity overcomes this repressive effect. To address this hypothesis, we first generated *Nkx2.1* knockout (KO) cells (*Nkx2.1*-KO; Supplementary Fig. S3A) and investigated their effects.

*Nkx2.1*-KO cells were differentiated for seven days without or with 5, 50 and 500 nM SAG, and Nkx2.2 expression was investigated using immunofluorescence. As a result, Nkx2.1-positive cells were no longer detected, confirming that *Nkx2.1* KO lines were successfully generated (Fig. 3A,C,E,F,H). With 5 nM SAG, a larger number of Nkx2.2-expressing cells were detected compared to wild-type (WT) cells, suggesting that Nkx2.1 has an essential function in repressing *Nkx2.2* repression (Fig. 3B,D,E). However, this aberrant increase was less evident at higher concentrations of SAG, and Nkx2.2 expression was rather perturbed in the *Nkx2.1*-KO cells at 500 nM SAG (Fig. 3E,G,I), suggesting that there is another mechanism by which Nkx2.2 expression is enhanced by Nkx2.1. Moreover, an increased population of Pax6-positive cells was detected at this concentration (Fig. 3J,K, Supplementary Fig. S4A), which corresponded to a decrease in the number of Nkx2.2-positive cells. This is consistent with the previous finding that Nkx2.2 and Pax6 are mutually repressive (Balaskas et al., 2012). There was no significant change in Rx expression (Fig. 3L,M, Supplementary Fig. S4B), suggesting that Rx is regulated independently of Nkx2.2 and Pax6 expression.

**Fig. 3.**
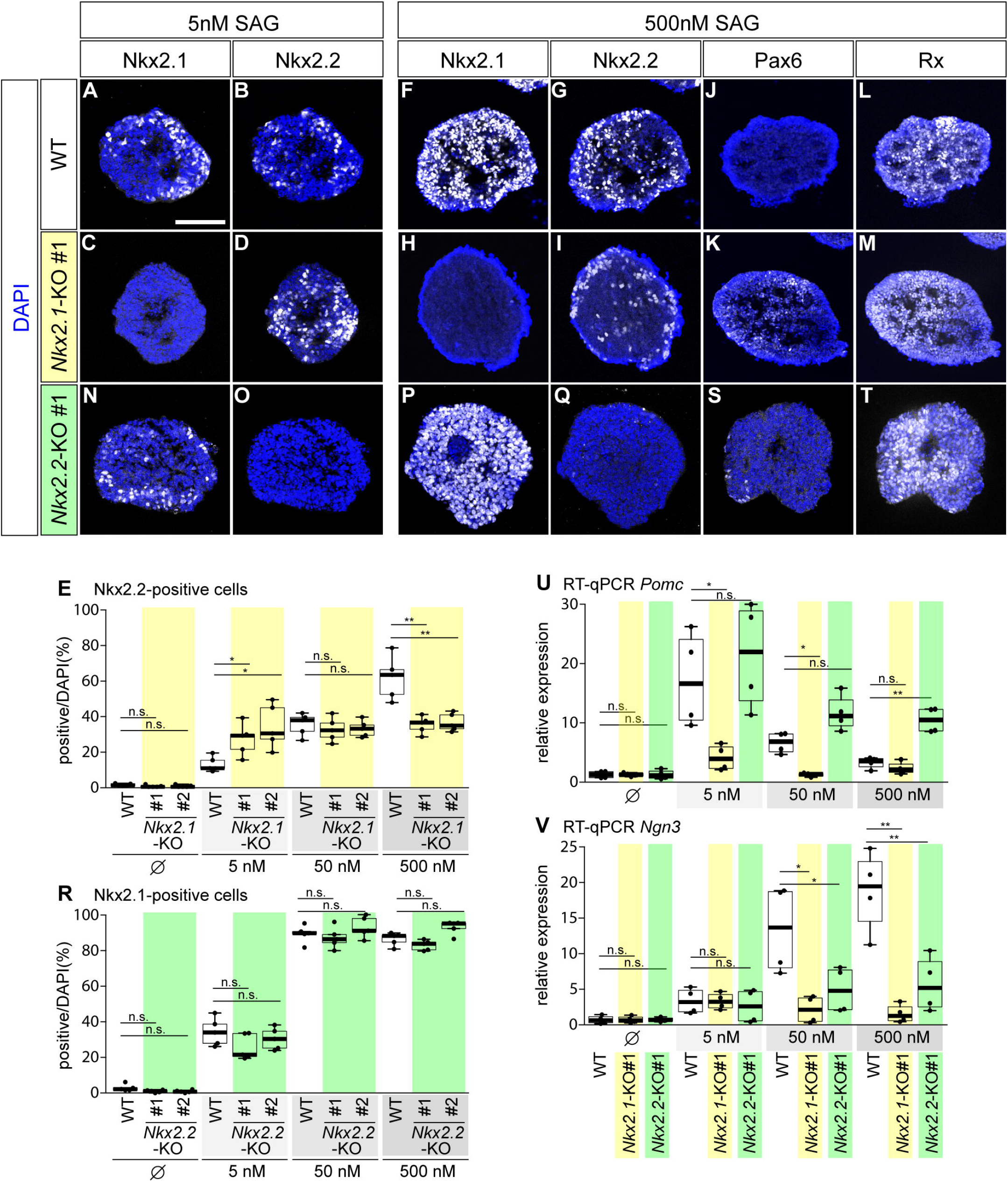
Nkx2.1 and Nkx2.2 are regulated in an epistatic manner. WT (A,B,F,G,J,L) *Nkx2.1*-KO (C,D,H,I,K,M) and *Nkx2.2*-KO (N,O,P,Q,S,T) cells were differentiated with 5 nM (A-D,N,O) or 500 nM SAG (F-M,P,Q,S,T) as in Fig. 2, and the colonies were analysed using immunofluorescence with antibodies against Nkx2.1 (A,C,F,H,N,P), Nkx2.2 (B,D,G,I,O,Q), Pax6 (J,K,S) and Rx (L,M,T). Scale bar, 50 m (A for A-D,F-Q,S,T). (E) Quantification of the positive cells for Nkx2.2 in WT and in the *Nkx2.1*-KO (yellow background) cells. (R) Quantification of the positive cells for Nkx2.1 in WT and in the *Nkx2.2*-KO (green background) cells. Note that the quantifications of WT are identical to those in Fig. 2Q,R,T,U. (U,V) Relative expression levels of *Pomc* (U) and *Ngn3* (V), as analysed by RT-qPCR. The expression data in WT are identical to those in Fig. 2V,W. The full expression data, together with that of *NR5A1*, is available in Supplementary Fig. S5.

Next, we asked whether Nkx2.2 counteracts Nkx2.1 expression. For this purpose, we further generated *Nkx2.2*-KO cells (Supplementary Fig.S3B) and analysed their effect on the expression of Nkx2.1 and Pax6. While Nkx2.2 was no longer detectable (Fig. 3O,Q,R), Nkx2.1 was expressed normally at all concentrations of SAG (Fig.3N,P,R), and Pax6 expressing cells were actually undetectable at 500 nM SAG (Fig. 3S, Supplementary Fig. S4A). Moreover, Rx expression was unaffected (Fig. 3L,T, Supplementary Fig. S4B). Thus, Nkx2.2 is dispensable for the initial expression of Nkx2.1 and repression of Pax6.

We further investigated neuronal subtypes differentiated in *Nkx2.1*- and *Nkx2.2*-KO cells. Each mutant cell was differentiated using different concentrations of SAG until day 12, as in the previous experiment (Fig. 2V–X). RT-qPCR revealed that the regional genes encoding the ARC marker *Pomc* and VMH markers *Ngn3* and *NR5A1* were not overall expressed in the *Nkx2.1*-KO cells (Fig. 3U,V, Supplementary Fig. S5A,C,E), suggesting that *Nkx2.1* is required for the differentiation of large subtypes of hypothalamic neurons. In contrast, *Nkx2.2*-KO cells exhibited downregulation of *Ngn3* and *NR5A1*, (Fig. 3V, Supplementary Fig. S5D,F) and aberrantly upregulated *Pomc* at 500 nM SAG (Fig. 3U, Supplementary Fig. S5B). This observation suggests that Nkx2.1 and Nkx2.2 are required for the proper differentiation of neuronal subtypes.

Taken together, these results show that *Nkx2.1* is epistatically upstream of *Nkx2.2* and governs Nkx2.2 expression both negatively and positively in a Hh signal-dependent manner.

### Nkx2.1 and Nkx2.2 are mutually repressive cell-autonomously, but are inducible non-cell autonomously

While Nkx2.1 and Nkx2.2 are mutually repressive at low concentrations of SAG, this effect is likely to be dependent on SAG concentration, and particularly Nkx2.2 expression requires Nkx2.1 at high SAG concentrations. To account for this complicated regulation, we next took advantage of the gene overexpression system in chick embryos and sought to determine the effect on each gene.

First, we investigated the expression of transcription factors during early developmental processes in the hypothalamus of chick embryos and found that the expression pattern was highly similar in mice and chick (Fig. 1, Supplementary Fig. S6), suggesting that the two species have similar developmental mechanisms.

Next, we overexpressed *control* (Fig. 2A–D’), *Nkx2.1* (Fig. 2E–H’) or *Nkx2.2* (Fig. 2I–L’) genes using electroporation at HH stage 8 (Hamburger and Hamilton, 1992) before the neural tube was closed and harvested 48 h post-electroporation (hpe), equivalent to HH stage 17.

To investigate the detailed patterning of the neural tube, cross-sections of the diencephalic area were prepared and the distributions of Nkx2.1 (Fig. 4A,A’,E,E’,I,I’), Nkx2.2 (Fig. 4B,B’,F,F’,J,J’) and Pax6 (Fig. 4C,C’,G,G’,K,K’) were analysed. Compared to that in *control*-electroporated embryos (*n* = 0/5 for all genes analysed), the expression of Nkx2.2 was suppressed in *Nkx2.1*-electroporated cells (Fig. 4F,F’; open arrowheads; *n* = 6/6 embryos). However, Nkx2.2-expressing cells appeared in the area adjacent to the electroporated cells (Fig. 4F,F’; filled arrowheads; *n* = 4/6 embryos), suggesting that Nkx2.1 induces the establishment of Nkx2.2-expressing area in surrounding cells, while Nkx2.1 represses Nkx2.2 expression in electroporated cells.

**Fig. 4.**
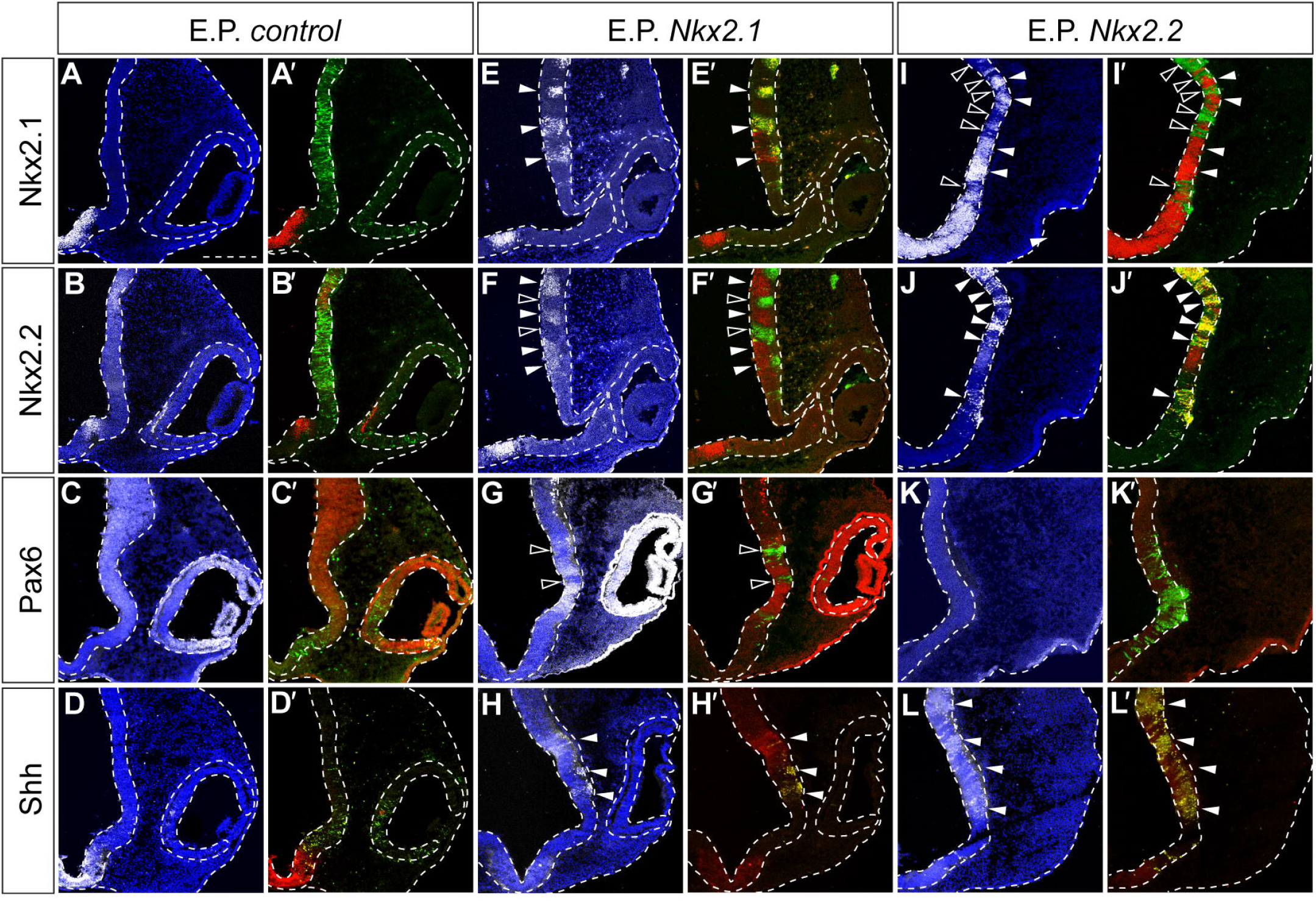
Nkx2.1 and Nkx2.2 repress mutual expression in cells but induce expression in surrounding cells. Expression plasmids harbouring *control GFP* (A–D’), *Nkx2.1* (E–H’) of *Nkx2.2* (I–L’) were electroporated at open neural plate area of head region at HH stage 8, and the embryos were analysed using immunofluorescence at 48 hpe. Hypothalamus area was analysed using antibodies against Nkx2.1 (white; A,E,I, red; A’,E’,I’), Nkx2.2 (white; B,F,J, red; B’,F’,J’), Pax6 (white; C,G,K, red; C’,G’,K’) and Shh (white; D,H,L, red; D’,H’,L’). Scale bar, 50 μm (A for all). Ectopically-induced expression and repressed expression are indicated by filled (E,E’,F,F’,H,H’,J,J’,L,L’) and open (F,F’,G,G’,I,I’) arrowheads, respectively. Scale bar, 50 μm (A for all).

Likewise, Nkx2.1 was downregulated in *Nkx2.2*-electroporated cells (Fig. 4I,I’; open arrowheads; *n* = 5/5 embryos); however, it was found to be upregulated in the adjacent cells (Fig. 4I,I’; filled arrowheads; *n* = 5/5 embryos). Thus, Nkx2.1 and Nkx2.2 are mutually repressive but induce each other’s expression in neighbouring cells.

We reasoned that this effect is mediated by the induction of a signal molecule, presumably Shh, by Nkx2.1 and Nkx2.2. To confirm this hypothesis, we performed immunostaining with an Shh antibody and found that Shh expression was upregulated in cells electroporated with Nkx2.1 (Fig. 4H,H’) and Nkx2.2 (Fig. 4L,L’’), suggesting that these two transcription factors induce Shh expression.

We also investigated *Shh* expression in hypothalamic neural cells differentiated from ES cells on day 7 (as in Fig. 2) using RT-qPCR. Increased *Shh* expression was observed upon treatment with SAG (Supplementary Fig. S1B). This observation suggests that hypothalamic cells induce *Shh* expression, and a positive feedback loop exists between the induction of Gli activity and *Shh* expression, which in turn induces Gli activity in the expressing cells and their surrounding cells.

Taken together, Nkx2.1 and Nkx2.2 are repressive, but non-cell-autonomously induce each other’s expression by mediating the induction of *Shh* expression.

### Rx is essential for positioning of hypothalamic region

Rx is also a transcription factor expressed in the anterior hypothalamus, and has been shown to be essential for its development (De Souza and Placzek, 2021; Orquera et al., 2016). Therefore, we attempted to understand how Rx is involved in hypothalamus development, especially in relation to the Shh-related genes *Nkx2.1* and *Nkx2.2*.

As neither did *Nkx2.1*- nor *Nkx2.2*-knockout cells exhibit any phenotypes for Rx expression (Fig. 3L,M,T), and Rx-positive cells were already found on day 4 before other hypothalamic transcription factors were expressed (i.e., Nkx2.1 and Nkx2.2; Supplementary Fig. S2D). We hence speculated that Rx may act upstream of these genes during hypothalamus development. To address this hypothesis, we used *Rx*-KO ES cells—where Rx function was lost—using CRISPR/Cas9 mutagenesis (Yamamoto et al., 2022) (Supplementary Fig. S3C). These mutant ES cells were normally maintained and differentiated into Pax6-positive neural cells without SAG (Fig. 5A–D) (Yamamoto et al., 2022), suggesting that early neural differentiation was normal in the Rx-mutant cells.

**Fig. 5.**
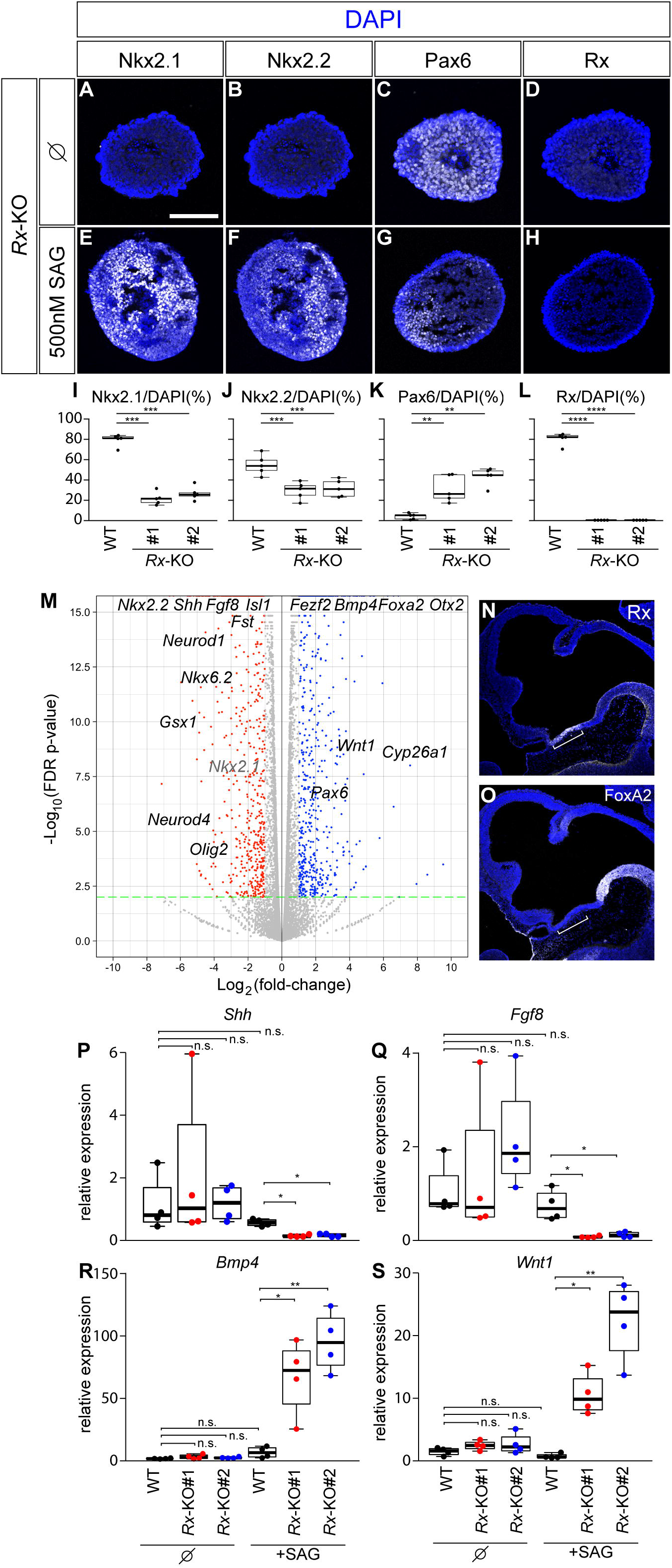
Rx is required for determination of hypothalamic area. (A–H) *Rx*-KO ES cells were differentiated with gfCDM in absence (A–D) or presence (E–H) of 500 nM SAG for seven days, and were sectioned and analysed using immunofluorescence with antibodies against Nkx2.1 (A,E), Nkx2.2 (B,F), Pax6 (C,G) or Rx (D,H). Scale bar, 50 m (A for A–H). (M) Cells deficient in Rx function fail to acquire hypothalamic identity. Volcano plot of high-throughput expression analysis on WT and *Rx*-KO cells differentiated using 500 nM SAG. Note that *Nkx2.1* (indicated with grey) is downregulated but not significantly (*p* > 0.05). (N,O) Rx and FoxA2 are expressed complementary to each other along the A-P axis. Immunofluorescence analysis of e10.5 sagittal sections was performed using Rx (N) and FoxA2 (O) antibodies. (P–S) RT-qPCR to evaluate the expression of *Shh* (P), *Fgf8* (Q), *Bmp4* (R) and *Wnt1* (S) in WT (black dots) or *Rx*-KO (red (for #1) and blue (for #2) dots) cells treated with 0 and 500 nM SAG.

However, an attempt to differentiate into hypothalamic cells with the same differentiation medium containing 500 nM SAG resulted in reduced numbers of Nkx2.1- and Nkx2.2-positive cells (Fig. 5E,F,I,J; compared to Fig. 2M,N) compared to those in WT cells (Fig. 2M,N,Q,R). Conversely, cells positive for Pax6 appeared abnormally (Fig. 5G,K).

These results suggest that Rx is required for pattern formation within the determination of the identities along the D-V axis during hypothalamus development; however, as Rx expression is not regulated by Shh and other hypothalamic genes, namely Nkx2.1 and Nkx2.2 (Fig. 2D,H,L,P,U and 3L,M,T), the regulation of D-V polarity by Rx remains unclear.

For an in-depth understanding of Rx function during hypothalamus development, we conducted high-throughput expression analysis based on RNA sequencing. RNAs were prepared from WT and *Rx*-mutant cells differentiated using 500 nM SAG for seven days (Fig. 2P).

Of the 25,683 mapped genes, 768 (indicated with blue dots; Fig. 5M) and 881 genes (indicated with red dots; Fig. 5M) were up and downregulated, respectively, in the *Rx*-mutant cells with more than 2-fold difference (*p* < 0.01 (Supplementary Tab. S1).

Notably, the expression of genes encoding signalling molecules that enable the identification of the rostral hypothalamic region was strongly dysregulated in *Rx*-KO cells. Enrichment was found for *Bmp4* and *Wnt1* in the KO cells, whereas decreased expression was detected for *Shh* and *Fgf8*. Given that *Shh* (Orquera et al., 2016) and *Fgf8* (Wexler and Geschwind, 2007) are expressed in the rostral diencephalic region, Rx is essential for the expression of genes enabling ventral hypothalamic identification. Moreover, *Bmp4* (Liu and Niswander, 2005) and *Wnt1* (Lavado et al., 2008; Martinez-Ferre and Martinez, 2009; Parr et al., 1993)—which are normally expressed in the dorsal diencephalon—were aberrantly upregulated, suggesting that Rx is required for the appropriate expression of the signal molecules that determine ventral diencephalic identity. In particular, as *Bmp4* (Liu and Niswander, 2005) and *Wnt1* (Lavado et al., 2008; Martinez-Ferre and Martinez, 2009; Parr et al., 1993) are expressed posteriorly to the hypothalamic region, the results suggest that A-P polarity was perturbed in *Rx*-KO cells (Acampora et al., 1997). Consistently, *Foxa2*—normally expressed posteriorly to Rx—(Fig. 5N,O) and *Otx2*—whose expression is complementary to that in the hypothalamus (Acampora et al., 1997)—were upregulated in *Rx*-KO cells. Furthermore, the neurogenic genes *Neurod1*, *Neurod4* and *Islet1* were downregulated (Fig. 5M), presumably because the expression of these signalling molecules was perturbed.

Next, we investigated the expression of signalling molecules in the absence and presence of SAG. As in the previous analysis (Fig. 2), both mRNAs from day 7 cells, either treated or untreated with 500 nM SAG, were examined using RT-qPCR. The expression of *Shh* (Fig. 5P), *FGF8* (Fig. 5Q), *Bmp4* (Fig. 5R) and *Wnt1* (Fig. 5S) was comparable between WT and *Rx*-KO cells in the absence of SAG. Upon treatment with SAG, WT cells expressed these genes at levels similar to those observed in the absence of SAG, suggesting that SAG did not alter the expression of these signalling molecules. However, the expression of these genes was greatly altered in *Rx*-KO cells. These observations suggest that the effects observed in *Rx*-KO cells occur only when hypothalamic differentiation is attempted.

Taken together, Rx is essential for the proper arrangement of the signal molecules that help identify the hypothalamic area along the D-V and A-P axes and act upstream of the gene regulatory networks for hypothalamus patterning.

### Hh signalling demarcates hypothalamus area from the retinal field

Although the two domains of the hypothalamus and retina were adjacent to each other, some of the hypothalamic cells were positive for Gli activity, while retinal cells were negative (Fig. 1C). Therefore, we inferred that hypothalamic and retinal cell identities were determined in a Gli-activity-dependent manner. To test this hypothesis, we disturbed Gli activity by overexpressing *Shh* and investigated the phenotype of the retina.

We found that the retinal area was severely reduced upon the overexpression of *Shh* (Fig. 6A–D; *n* = 5/8 embryos, and 3/8 embryos dwarfed). Sectional analysis further revealed an increase in Nkx2.1- (Fig. 6E,E’; in comparison with Fig. 4A,A’ and Supplementary Fig. S6E) and Nkx2.2- (Fig. 6F,F’; Fig. 4B,B’, Supplementary Fig. S6F)-expressing cells (*n* = 6/6) and complete inhibition of Pax6 expression (Fig. 6G,G’; Fig. 4C,C’, Supplementary Fig. S6G; *n* = 6/6). However, Rx expression was less evidently altered by the overexpression of *Shh* (Fig. 6H,H’; Supplementary Fig. S6H; *n* = 6), suggesting that Hh signalling induces ventral hypothalamic identity.

**Fig. 6.**
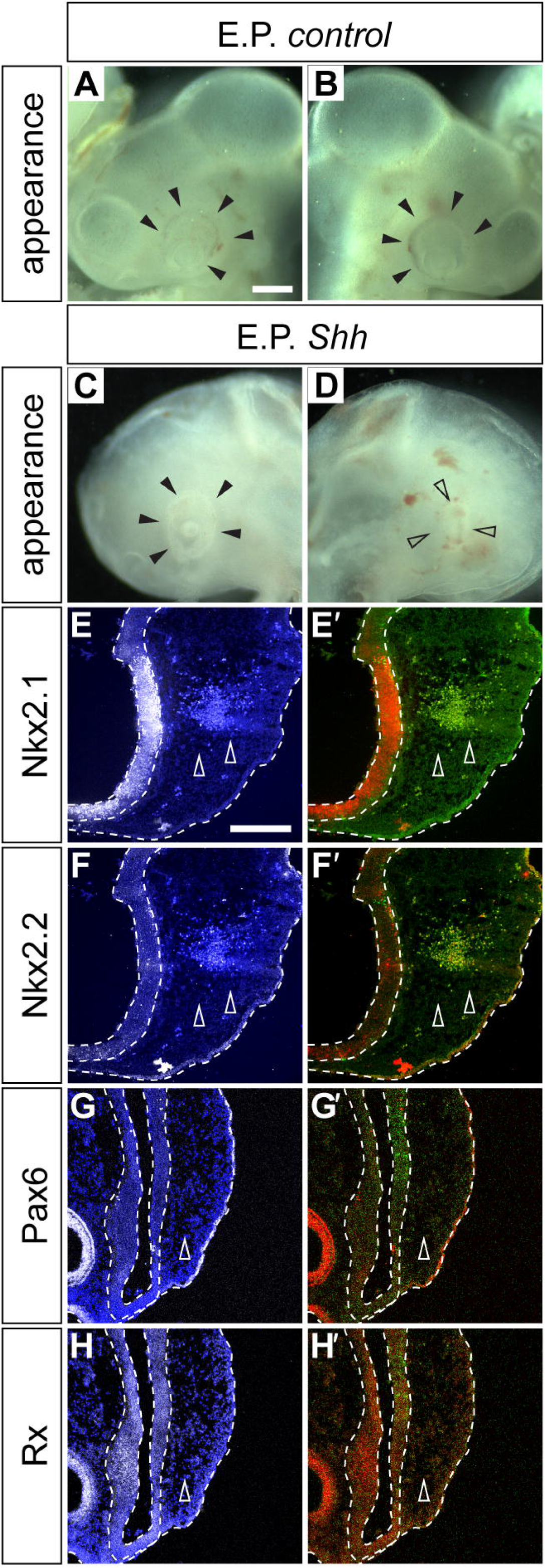
Shh demarcates hypothalamic region from retinal area in ventral diencephalon. Head appearance of embryos electroporated with *control* (A,B; as in Fig. 4A–D’) or *Shh* (C,D). Eye regions are indicated by black and open arrowheads. Shh expression plasmid was electroporated, and the embryos were analysed as shown in Fig. 4. (A,B) Appearance of head regions of embryos electroporated with *Shh* expression plasmid, from right (A) and left (B) sides. (C–H’) Phenotypes observed upon transfection of expression vector harbouring *Shh* were analysed using Nkx2.1 (E,E’), Nkx2.2 (F,F’), Pax6 (G,G’) and Rx (H,H’). Scale bar, 200 μm (A for A–D), and 100 μm (E for E–H’).

These data demonstrate that Shh signalling separates the retinal area from the hypothalamic area during the development of the ventral diencephalon.

## Discussion

### The differentiation system of mutant ES cells provides direct information about the relationship between signal inputs and gene expression

In the present study, we demonstrated that the subdomains of the hypothalamus were patterned in a Gli-activity-dependent manner, and we recapitulated this patterning using the differentiation system from ES cells (Fig. 1, 2). Moreover, we showed that mutual transcriptional regulation was required for the expression of these hypothalamic transcription factors using a series of CRISPR/Cas9-generated mutants (Fig. 3). Gene overexpression analysis revealed that the hypothalamic transcription factors demarcated the neighbouring retinal area (Fig. 4, 6).

The importance of morphogens in patterning has been suggested in several organs, particularly those comprising epithelial cells (e.g., the nervous system) (Christian, 2012; Sagner and Briscoe, 2017). Signalling gradients induce the expression of different sets of genes; their combinations confer cellular diversity and cells are deposited in specific proportions. KO mouse models are effective with respect to proving the critical roles of these genes in the precise patterning of organs. However, caution is warranted in this approach as genes are often expressed in a number of different areas from the one in focus, and the observed phenotypes do not exclude the possibility that the observed phenotypes could be a secondary outcome caused by their influence on different areas. This is the case for the transcription factors investigated in the current study, where they are expressed in a variety of brain regions.

In addition, the simple gene knockout approach has a possible problem in which the lineage of the cells in the WT and KO individuals cannot be directly compared. The conditional KO strategy, in which gene expression is attenuated in a specific area, is a good experimental system to circumvent these limitations (Skarnes et al., 2011). The approaches taken in the present study, in which we differentiated mutant ES cells directly into hypothalamic cells, also allowed us to evaluate the direct effect of the attenuation of a particular gene while excluding the potential of a secondary effect.

### Hh signal triggers transcriptional networks and induces diversity in hypothalamic cell types

Signal gradients are important for patterning a wide spectrum of organs (Sagner and Briscoe, 2017). The expression of *Nkx2.1* and *Nkx2.2* was induced by different intensities of Gli activity (Fig. 1) or SAG concentrations (Fig. 2), showing that morphogens used in the ventral diencephalic area confer distinct ventral (positive for Nkx2.1) and dorsal (double positive for Nkx2.1 and Nkx2.2) hypothalamic identities in the forebrain progenitor cells (Fig. 7).

**Fig. 7.**
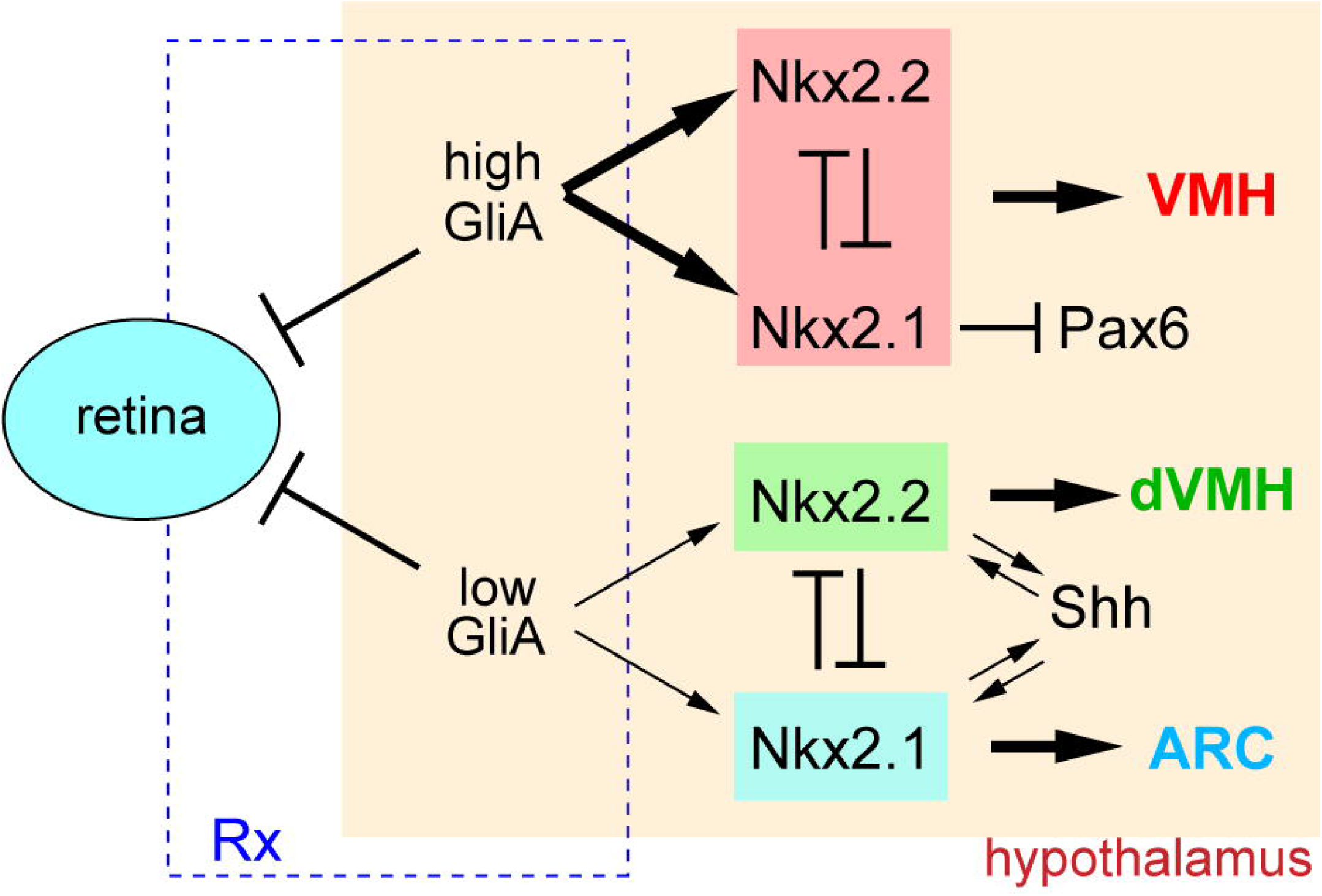
Working hypothesis of transcriptional network leading to multiple cell types by Shh-related genes. At low-Gli activity (low GliA), either Nkx2.1 or Nkx2.2-positive cells appear separately and a mutual repression is observed between these molecules. High-Gli activity (high GliA) overrides the repressive effects of Nkx2.1 and Nkx2.2, leading to the expression of both genes. Different combinations of transcription factors lead to differentiation, resulting in distinct hypothalamic identities. Gli activity triggered by Shh blocks retinal development. Rx acts upstream of Nkx2.1 and Nkx2.2 and is required for the development of hypothalamus as well as the retina (depicted as a rectangle with dashed lines). Shh expression is induced by both Nkx2.1 and Nkx2.2, and Pax6 expression is repressed by Nkx2.1. VMH; ventromedial hypothalamus, dVMH; dorsal part in VMH region, ARC; arcuate nucleus.

Nkx2.2 and Pax6 are in a repressive relationship in the spinal cord (Balaskas et al., 2012; Briscoe et al., 1999). Our data suggest that this is the case in the ventral diencephalon, where Nkx2.2 and Pax6 are expressed in a complementary manner (Fig. 1L,O,R,U) and are mutually repressive (Fig. 2B,C,F,G,J,K,N,O). Moreover, Nkx2.2 expression showed increased sensitivity to Hh (Fig. 3B,F,J,N,R). In addition, overexpression of Nkx2.1, blocked Pax6 expression (Fig. 4G,G’). Therefore, it is likely that Nkx2.1 expression induced by Hh signalling, gradually repressed Pax6 expression, allowing the initiation of Nkx2.2 expression. This is similar to the network formed by three transcription factors Olig2, Nkx2.2 and Pax6 in the ventral spinal cord; Olig2 expression is induced by Hh, and Olig2 represses Pax6 expression, which triggers Nkx2.2 expression (Balaskas et al., 2012).

While Nkx2.1 and Nkx2.2 repress each other, they induce *Shh* expression (Fig. 4), suggesting that both act as transcription factors in a dual mode, repressors of *Nkx2.2/Nkx2.1* and activators of *Shh*. Nkx2.1 has been shown to function as both a repressor and an activator (Blaess et al., 2014), and Nkx2.2 can also function as a transcriptional activator (Watada et al., 2000) as well as a repressor (Kutejova et al., 2016). These findings are in agreement with our findings.

In the current study, we focused on three transcription factors (Nkx2.1, Nkx.2.2 and Pax6) whose expression is regulated by Hh. However, many other transcription factors are expressed in the same region as Nkx2.1 (Kim et al., 2020; Shimogori et al., 2010), and several of them may be the target genes of Hh. This suggests that transcriptional networks other than those between Nkx2.1, Nkx2.2 and Pax6 exist and provide more subdomains in the hypothalamus. To discover such subdomains and the transcriptional networks leading to them, it would be informative to identify the transcription factors induced by different concentrations of SAG through transcriptome analysis and to classify the genes with different dependencies on Hh signalling.

### The contribution of Rx to hypothalamus development and pattern formation

Rx is expressed in the hypothalamus and optic regions, but the regulation of its expression and its roles in development and pattern formation are more complicated than those of Nkx2.1 and Nkx2.2, whose expression is directly induced by Hh (Kutejova et al., 2016; Vokes et al., 2007). Rx begins to be expressed earlier than Nkx2.1 and Nkx2.2 (Supplementary Fig. S2); thus, the Rx acts upstream of the hypothalamic transcription factors and provides the basis for the ventral diencephalic region.

Rx is expressed only in the forebrain and later in diencephalic areas, but not in the spinal cord (Furukawa et al., 1997; Mathers et al., 1997). This restricted expression with respect to the A-P axis is similar to another homeodomain-containing transcription factor, Otx2, which is expressed only in the forebrain and midbrain (Kurokawa et al., 2014). Therefore, the Rx demarcates the hypothalamic region not only along the D-V axis but also along the A-P axis.

We showed that Rx regulates the expression of signalling factors that indicate the forebrain/diencephalic area (Fig. 5M). It has also been shown that another homeobox transcription factor, Six3, has an expression pattern and activity similar to that of Rx; Six3 is expressed in the hypothalamus and optic vesicle (Oliver et al., 1995), and embryos devoid of Six3 exhibited aberrant *Wnt1* induction (Lagutin et al., 2003), caudalisation of the forebrain region (Lagutin et al., 2003; Lavado et al., 2008) and failure to activate *Shh* expression (Geng et al., 2008), suggesting that Six3 and Rx share functions in terms of the spatial assignment of the diencephalon. However, Six3 expression does not depend on Rx, as revealed by the expression profiling (Fig. 5M; Supplementary Tab. S1), suggesting that a strong regulation does not exist between these two factors. Chromatin immunoprecipitation sequencing (ChIP-seq) is useful for identifying their binding sites and direct target genes.

Overall, the present study has revealed a mechanism in which a combination of permissive (i.e., Rx expression) and inductive effects (i.e., gradient of Shh) is important for the spatiotemporally precise assignment of hypothalamic identities. Rx confers the ventral diencephalic field, thereby providing multiple types of hypothalamic progenitor cells that are positioned in an orderly manner.

Future analyses are important for identifying downstream transcriptional networks and secondary signals that promote neuronal differentiation. By correlating each early hypothalamic subdomain with its functional component, we will be able to trace cell lineages for the diverse functions of the hypothalamus.

## Materials and Methods

### Ethics statement and *Tg (GBS-GFP)* transgenic mice

All animal experiments were performed with the approval of the Animal Welfare and Ethics Committee of the Nara Institute of Science and Technology (approval number: 1810 (for mice), 1636 (for chick) and 389 (for genetic modifications)) and were in accordance with the relevant guidelines and regulations.

The transgenic *Tg (GBS-GFP)* mouse line was kindly provided by Dr. James Briscoe. In these mice, GFP expression was driven by octameric Gli-binding sites derived from the FoxA2 gene enhancer (Balaskas et al., 2012). Breeding pairs were set up, and the noon of the day when vaginal plug was confirmed was assumed to be embryonic day of 0.5.

### Maintenance and differentiation of mouse ES cells

Sox1-GFP murine ES cells (Ying et al., 2003) were maintained in medium containing double inhibitors (Ying et al., 2008). Differentiation into hypothalamic cells was performed according to an established protocol (Wataya et al., 2008), with the exception that SAG—an agonist for smoothened (Selleck; S7779)—was used instead of recombinant Shh protein.

For differentiation, three thousand ES cells were seeded onto a 96-well non-absorbable plate (Greiner, Cell-Repellent, #650970) and cultured in growth factor-free chemically defined medium (gfCDM) containing Iscove’s Modified Dulbecco’s Medium (IMDM; SIGMA #I3390) and Ham’s F-12 (Wako, #087-08335) at a 1:1 ratio, supplemented with chemically defined lipid concentrate (Thermo #11905031), Penicillin-Streptomycin-L-Glutamine (Wako, #161-23201), 0.2% Bovine Serum Albumin (SIGMA, #A9418), and 450 nM α-monothioglycerol (MP Biomedicals, #02155723-CF).

We did not observe any Nkx2.1- or Nkx2.2-positive cells in the absence of SAG, although it has been shown that such ventral hypothalamic cells appear regardless of the absence of SAG (Kodani et al., 2022; Wataya et al., 2008). This is presumably due to the differences of the cell lines used in each study.

### Generation of gene-modified ES cells

Gene-modified ES cells were generated via CRISPR/Cas9 mutagenesis. Guide RNAs targeting *Nkx2.1* (5′-CCATGCAGCAGCACGCCGTG-3′), *Nkx2.2* (5′-AAGACGGCTCGGTGGCCGAA-3′) and *Rx* (5′-GGAGCCTCCGGCTGGCGCCT-3′) were identified using the CRISPR website (https://crispr.dbcls.jp), and the synthesised DNA fragments were subcloned into the BbsI sites of the *pX459* vector (#62988, Addgene). The resulting constructs were transfected into ES cells using Lipofectamine 3000 (Life Technologies, #L3000001). Transfected cells were selected via puromycin selection (1 μg/mL for 2 days). Two clones in which the mutation had been introduced homologously were selected from each gene knockout, and the mutations were verified by sequencing (Supplementary Fig. S3). Attenuated expression was further verified using immunofluorescence for proteins encoded by the target genes. One of the *Rx*-KO clones used in the present study was used in our previous study (Supplementary Fig. S3C; #1) (Yamamoto et al., 2022), and another clone was used to verify this phenotype (Fig. 5, Supplementary Fig. S3C; #2).

### Electroporation of chick embryos

Fertilised chicken eggs were purchased from Yamagishi farm (Wakayama Prefecture, Japan), and the developmental stage was evaluated in accordance with the Hamburger and Hamilton criteria (Hamburger and Hamilton, 1992). Electroporation was performed using an electroporator (BTX830; five pulses at 25 V, each of which was 50 ms, at 450 ms intervals). Embryos were harvested 48 h after electroporation and subjected to further analysis.

### Histological analysis and quantification

For histological analyses, the harvested embryos or colonies were fixed in 4% paraformaldehyde for one hour and subsequently incubated overnight at 4°C with 15% (w/v) sucrose in a rotating incubator. The specimens were embedded in O.C.T. compound (Sakura Finetek) and cryosectioned at a thickness of 12 μm on a Polar cryostat (Sakura Finetek). For immunofluorescence, the following antibodies were used: Rx (rabbit; TaKaRa, #M228; 1:1,000), Nkx2.1 (rabbit; Abcam, #ab76013; 1:1,000), Pax6 (rabbit; Millipore, #AB2237; 1:1,000, and mouse; Developmental Studies Hybridoma Bank [DSHB], #PAX6; 1:50), Nkx2.2 (mouse; DSHB, #74.5A5; 1:50), Shh (mouse; DSHB, #5E1; 1:50) and GFP (sheep; AbD Serotec, #4745-1051). The secondary antibodies used were rabbit IgG (Jackson Laboratories, #711-166-152 for Cy3; #711-606-152 for Cy5), mouse (#715-166-151 for Cy3; #715-606-150 for Cy5) and sheep (#713-096-147 for FITC), all at 1:500.

For differentiation of ES cells, eight colonies were prepared and fixed, and sections were prepared for each differentiation condition. After immunofluorescence, five samples were randomly chosen, and positive cells for each gene were counted on sectional planes where the largest number of DAPI-positive cells in each colony was found (containing approximately 300–400 cells).

### RNA extraction and RT-qPCR

Total RNA extraction and reverse transcription were performed using NucleoSpin (Takara, #U955C) or Picopure (Thermo, # KIT0204) RNA extraction kits, and PrimeScript™ RT Master Mix (Takara, RR036), respectively. qPCR was performed on a CFX qPCR machine (Bio-Rad) using the primers listed in Supplementary Tab. S2. Amplification data were analysed using the comparative C_t_ method and target gene expression was normalised to that of *RhoA*.

### High-throughput expression analysis

WT and *Rx*-mutant cells were differentiated using the hypothalamus differentiation protocol (with 500 nM SAG from days 3–7), and RNA was extracted. The libraries were prepared using the TruSeq stranded-mRNA library preparation kit (Illumina, #20020594) and sequenced using the NextSeq 500 platform (Illumina). Approximately 20 million reads were obtained per sample and mapped with the reference sequences of 25,683 genes using CLC genomics workbench software (Robinson et al., 2010).

### Image processing and statistics

Images were captured using LSM710 and LSM980 confocal microscopes (Zeiss, Germany) and processed using the Photoshop software (Adobe). Figures were prepared using Illustrator (Adobe) and box plots were created using R software (version 4.2.1). Differences were evaluated using either a two-tailed Student’s *t-*test (for comparisons between two groups) or one-way analysis of variance (ANOVA; for comparisons between more than two groups). Statistical comparisons with *p* < 0.05 were considered significant. *p*-values (\**p* < 0.05; ** *p* < 0.01; *** *p* < 0.001; **** *p* < 0.0001) are indicated in each graph.

## Data availability

Raw RNA-sequencing data are available from the DNA Data Bank of Japan (DDBJ, https://www.ddbj.nig.ac.jp/index-e.html) with accession number PRJDB13538.

## Acknowledgements

The authors thank James Briscoe for the *Tg (GBS-Gfp)* line, Kazuaki Takahashi for technical assistance, Takaaki Matsui for comments, and all other laboratory members, particularly Saori Yoshinaga, for their support and discussion. This work was supported in part by a grants-in-aid for scientific research from the Japan Society for the Promotion of Science (19H04781 and 20H03263) to NS.

## Competing interests

The authors declare no competing interests exist.

**Supplementary materials can be found as a separate file.**

## Notes

### Competing Interest Statement

The authors have declared no competing interest.

